# Venomics AI: a computational exploration of global venoms for antibiotic discovery

**DOI:** 10.1101/2024.12.17.628923

**Authors:** Changge Guan, Marcelo D. T. Torres, Sufen Li, Cesar de la Fuente-Nunez

## Abstract

The relentless emergence of antibiotic-resistant pathogens, particularly Gram-negative bacteria, highlights the urgent need for novel therapeutic interventions. Drug-resistant infections account for approximately 5 million deaths annually, yet the antibiotic development pipeline has largely stagnated. Venoms, representing a remarkably diverse reservoir of bioactive molecules, remain an underexploited source of potential antimicrobials. Venom-derived peptides, in particular, hold promise for antibiotic discovery due to their evolutionary diversity and unique pharmacological profiles. In this study, we mined comprehensive global venomics datasets to identify new antimicrobial candidates. Using machine learning, we explored 16,123 venom proteins, generating 40,626,260 venom-encrypted peptides (VEPs). Using APEX, a deep learning model combining a peptide-sequence encoder with neural networks for antimicrobial activity prediction, we identified 386 VEPs structurally and functionally distinct from known antimicrobial peptides. Our analyses showed that these VEPs possess high net charge and elevated hydrophobicity, characteristics conducive to bacterial membrane disruption. Structural studies revealed considerable conformational flexibility, with many VEPs transitioning to α-helical conformations in membrane-mimicking environments, indicative of their antimicrobial potential. Of the 58 VEPs selected for experimental validation, 53 displayed potent antimicrobial activity. Mechanistic assays indicated that VEPs primarily exert their effects through bacterial membrane depolarization, mirroring AMP-like mechanisms. *In vivo* studies using a mouse model of *Acinetobacter baumannii* infection demonstrated that lead VEPs significantly reduced bacterial burdens without notable toxicity. This study highlights the value of venoms as a resource for new antibiotics. By integrating computational approaches and experimental validation, venom-derived peptides emerge as promising candidates to combat the global challenge of antibiotic resistance.

## Introduction

Drug-resistant infections account for approximately 5 million deaths annually worldwide^1^, fueled by the rapid emergence of antibiotic-resistant pathogens. Among these, Gram-negative bacteria, identified as priority pathogens by the World Health Organization (WHO), are particularly adept at developing resistance. Despite this growing threat, the development pipeline for novel antibiotics has stagnated over the past few decades due to high costs and lengthy timelines, emphasizing the urgent need for innovative therapeutic strategies ^2,3^.

Animal venoms represent a promising source of new antibiotics^4–7^. These venoms are rich in bioactive peptides and proteins that exhibit diverse pharmacological effects, including antibacterial activity. Venom-derived peptides can target ion channels, cell membranes, and enzymes. For example, the cone snail toxin MVIIA (Ziconotide), marketed as Prialt®, is used to treat chronic pain by selectively targeting voltage-gated calcium channels^8^.

Evolutionary studies have shown that venom genes originated from a small number of ancestral genes and diversified rapidly, resulting in a vast reservoir of chemical diversity^5^.

Recent advances in bioinformatics and machine learning have enabled the systematic mining of potential antibiotic candidates from proteomes^9–16^. Using APEX, our sequence-to-function predictor^9,17^, here we systematically identified potential antibiotics within venom proteomes and experimentally validated their antimicrobial activity. Notably, we identified venom protein-derived encrypted peptides (VEPs) with antimicrobial efficacy both *in vitro* and in preclinical animal models **(Figure 1a).** These findings highlight the immense, untapped potential of venomics to address the global challenge of antibiotic resistance.

**Figure 1.**
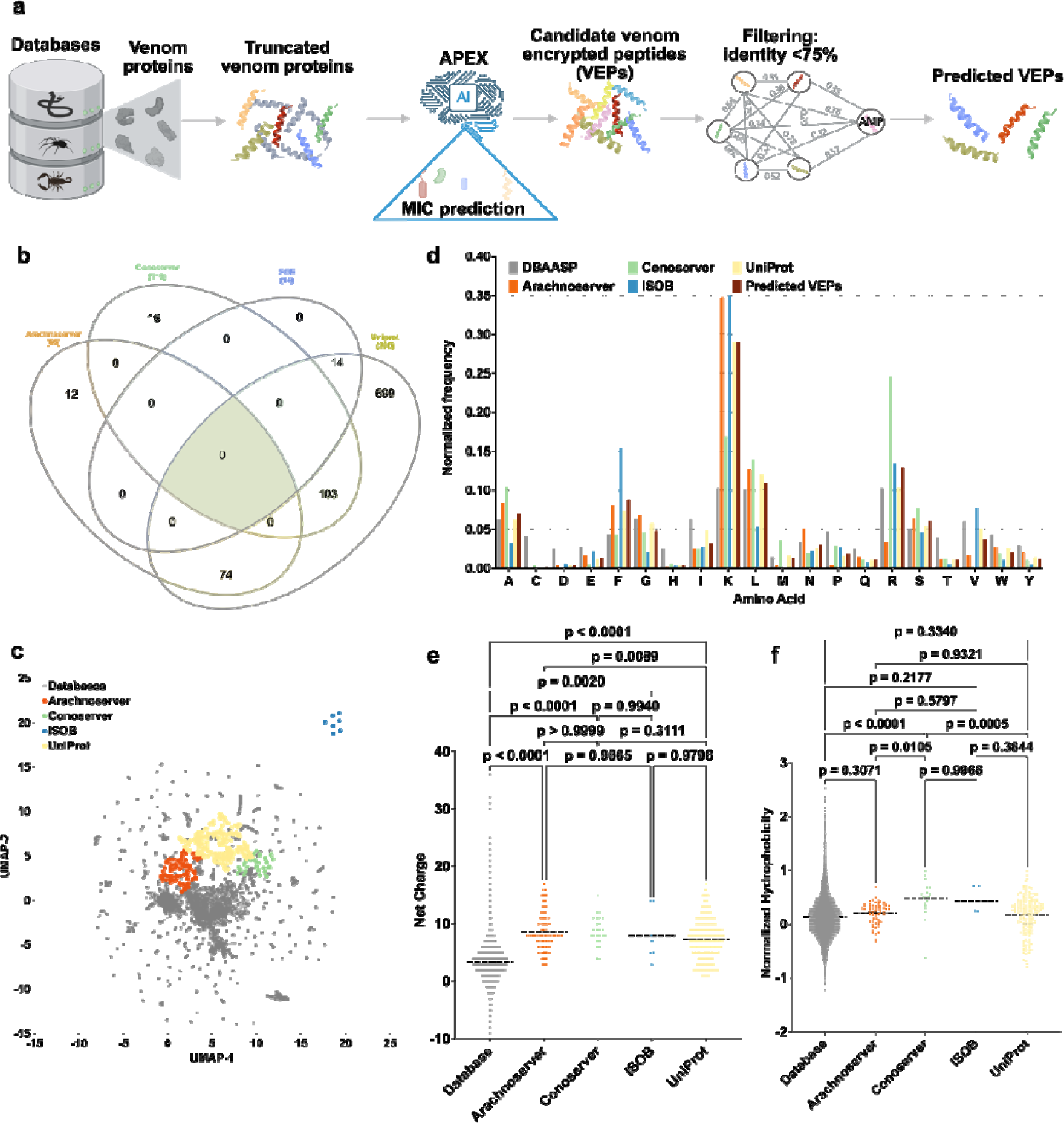
Exploration of global venoms for antibiotic discovery. **(a)** Mining framework for AMPs. Our framework employs a three-stage approach to identify novel AMP candidates from venom proteins. Initially, a peptide library is generated using a sliding window method, extracting peptides ranging from 8 to 50 amino acid residues in length. Subsequently, Minimum Inhibitory Concentration (MIC) values of a peptides against bacterial strains were predicted by APEX. Finally, candidate VEPs are selected based on sequence similarity, yielding a set of unique and potent molecules. **(b)** Venn diagram illustrating species overlap among the four databases used as source of venom proteins. Species names extracted from these databases were analyzed to identify diversity. **(c)** Physicochemical feature space exploration. The graph illustrates a bidimensional sequence space visualization of peptide sequences found in DBAASP and antimicrobial venom-derived EPs (VEPs) discovered by APEX in venom proteins from multiple source organisms. The physicochemical features were calculated for peptide sequences which was made up of the feature vector for representing peptide. Each row in the matrix represents a feature representation of a peptide based on its amino acid composition. Uniform Manifold Approximation and Projection (UMAP) was applied to reduce the feature representation to two dimensions for visualization. **(d)** Comparison of amino acid frequency in VEPs with known antimicrobial peptides (AMPs) from the DBAASP, APD3, and DRAMP 3.0 databases. Distribution of two physicochemical properties for peptides with predicted antimicrobial activity, compared with AMPs from DBAASP, APD3, and DRAMP 3.0: **(e)** net charge and **(f)** normalized hydrophobicity. Net charge influences the initial electrostatic interactions between the peptide and negatively charged bacterial membranes, while hydrophobicity affects interactions with lipids in the membrane bilayers. Statistical significance in **e** and **f** was determined using two-tailed t-tests followed by Mann–Whitney test; P values are shown in the graph. The solid line inside each box represents the mean value for each group.

## Results

### Mining venoms for new antibiotics

We sourced venom proteins from four databases: ConoServer (focusing on conopeptides, **Supplementary Figure 1**),^18^ ArachnoServer (spider proteins, **Supplementary Figure 2**), ^20^ ISOB (indigenous snake proteins, **Supplementary Figure 3**),^22^ and VenomZone (covering six taxa: snakes, scorpions, spiders, cone snails, sea anemones, and insects, **Supplementary Figure 4**)^24^. The VenomZone dataset, curated from UniProtKB, was represented in our study by UniProt. Altogether, we compiled 16,123 venom proteins, which were computationally processed to generate 40,626,260 VEPs.

To analyze differences across the four databases, we performed a species overlap analysis (**Figure 1b**). UniProt contained the largest number of unique species (699), reflecting its extensive coverage. Conoserver and Arachnoserver encompassed smaller unique subsets (16 and 12, respectively), while ISOB contained no unique species. These results highlight the complementary nature of these databases, emphasizing the value of integrating multiple sources to achieve comprehensive venom protein diversity.

Using APEX, a deep learning model, we predicted bacterial strain-specific MIC values for each peptide and used the mean MIC as a measure of antimicrobial activity. We identified 7,379 VEPs with a mean MIC ≤32 µmol L^-1^ (**Data S1**). Further filtering criteria (see **Methods**: “**Venom encrypted peptide selection**”) based on sequence similarity to known antimicrobial peptides (AMPs) yielded 386 candidates with low similarity to existing AMPs (**Supplementary Table 1** and **Data S2**).

To visualize sequence diversity, we compared the 386 VEPs with 19,762 known AMPs from the DBAASP database. Pairwise alignment (see **Methods**: **“Peptide sequence similarity”**) and uniform manifold approximation and projection (UMAP) revealed that most known AMPs clustered densely, reflecting high sequence similarity matrix (**Figure 1c**).

Most known AMPs formed a dense cluster, indicating high sequence similarity, with a minority scattered outside this cluster, representing more diverse sequences. VEPs derived from ConoServer and ArachnoServer tended to cluster closer to known AMPs, reflecting relatively higher sequence similarity. In contrast, UniProt-derived VEPs mapped farther from the AMP cluster, partially overlapping with scattered AMPs and occupying previously unexplored regions of sequence space. ISOB-derived VEPs were the most distant from known AMPs, forming isolated clusters that represent a promising source of completely novel AMP sequences (**Figure 1c**).

To determine whether VEPs with low sequence similarity to known AMPs share key physicochemical characteristics, we analyzed their distribution in physicochemical feature space (**Supplementary Figure 5**). While known AMPs from DBAASP clustered centrally, UniProt-derived VEPs formed three distinct clusters, Arachnoserver-derived VEPs formed two clusters, and ISOB and Conoserver each formed one cluster. UniProt cluster overlapped with ConoServer, while the ISOB-derived cluster remained entirely isolated. UniProt- and Arachnoserver-derived clusters that did not overlap with known AMPs represent unexplored regions of sequence space (**Figure 1c**).

These findings suggest that our approach identifies both AMP-like peptides that differ in sequence while sharing similar physicochemical properties and entirely different AMP families that deviate in both sequence and characteristics.

### Composition and physicochemical features

A comparison of amino acid composition between VEPs and DBAASP AMPs revealed distinct profiles (**Figure 1d** and **Supplementary Figure 6**). VEPs had lower cysteine, aspartic acid, histidine, and isoleucine, while showing higher phenylalanine, lysine, and arginine content. ISOB-derived VEPs were particularly enriched in phenylalanine, whereas Conoserver-derived VEPs displayed pronounced arginine content. Notably, Arachnoserver- and ISOB-derived VEPs were enriched in lysine.

To further understand the physicochemical properties contributing to antimicrobial activity, we benchmarked VEPs against known AMPs (**Figure 1e-f, Supplementary Figure 7**). VEPs were generally more positively charged, facilitating electrostatic interactions with the negatively charged bacterial membranes²². They also exhibited slightly higher normalized hydrophobicity, likely driven by their increased phenylalanine and arginine content. In ISOB- and Conoserver-derived VEPs, these features enhanced amphiphilicity (**Supplementary Figure 7a**), promoting secondary structure formation and membrane-associated activity.

Additionally, VEPs displayed higher isoelectric points than known AMPs (**Supplementary Figure 7b**), consistent with their elevated cationic residue content. By design, the APEX model excluded peptides with high cysteine content, thereby avoiding many Conoserver-derived peptides rich in disulfide bridges. Despite their elevated phenylalanine levels, VEPs maintained comparable normalized hydrophobic moments (**Supplementary Figure 7c**) and aggregation propensities (**Supplementary Figure 7d**) to conventional AMPs, with amphiphilic distribution likely mitigating hydrophobic clustering.

Collectively, these results delineate the unique composition and physicochemical properties of VEPs, highlighting their potential as promising antimicrobial candidates.

### Antimicrobial Activity Assays

Among the 58 VEPs tested, 53 (91.4%) exhibited activity against at least one pathogenic strain. Notably, all Arachnoserver-derived peptides were active, emphasizing their strong antimicrobial potential (**Figure 2a**). In contrast, some UniProt-derived VEPs (from VenomZone) demonstrated limited potency: UniprotKB-2 showed no activity, while UniprotKB-6 and UniprotKB-11 were active only against *Enterococcus faecium*.

**Figure 2.**
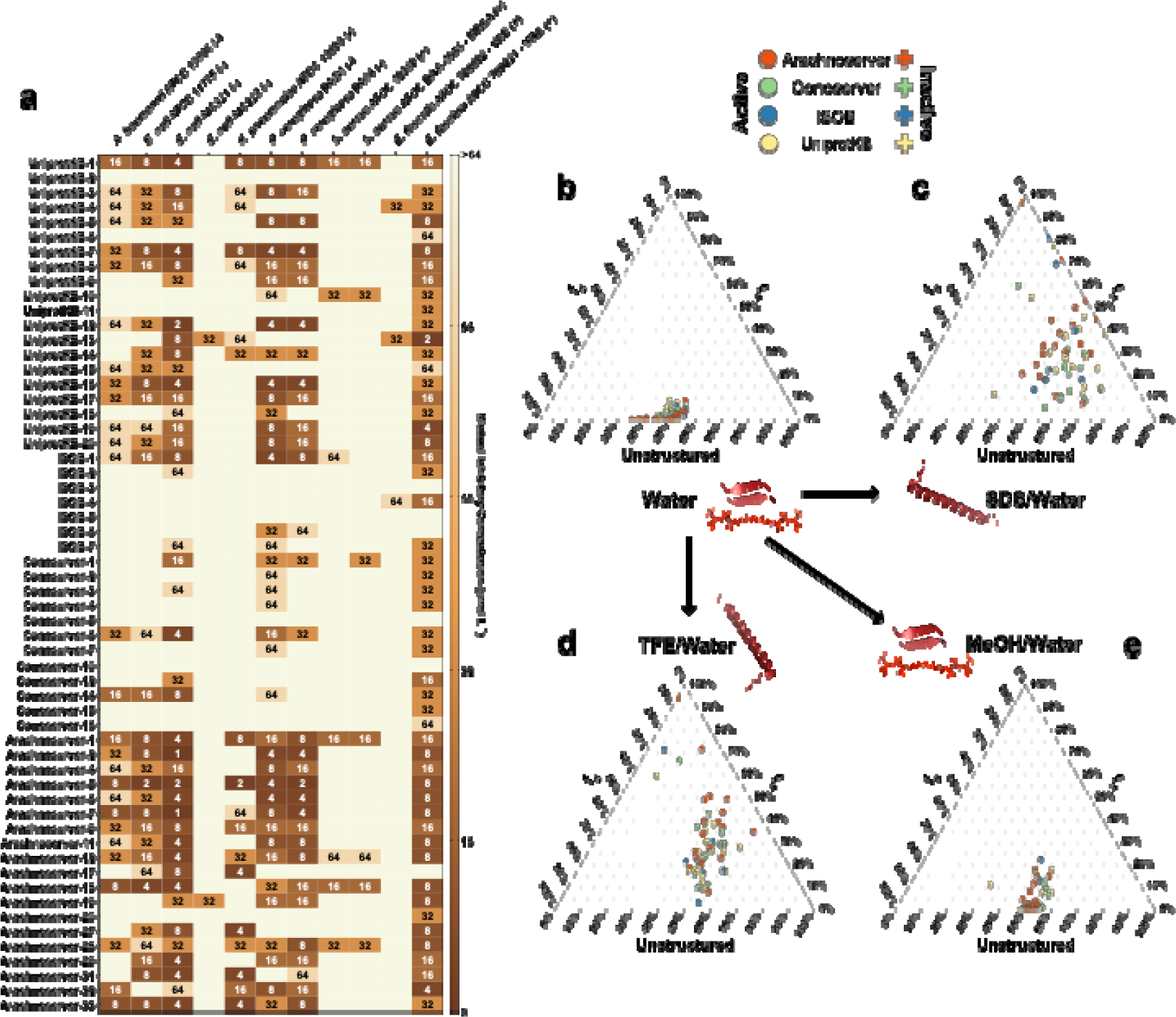
Antimicrobial activity and secondary structure profiles of antibiotics from venoms. **(a)** Heat map displaying the antimicrobial activities (μmolCL^-1^) of active antimicrobial agents from venoms against 11 clinically relevant pathogens, including antibiotic-resistant strains. Briefly, 10^5^ bacterial cells were incubated with serially diluted VEPs (1–64CμmolCL^-1^) at 37C°C. Bacterial growth was assessed by measuring the optical density at 600 nm in a microplate reader one day post-treatment. The MIC values presented in the heat map represent the mode of the replicates for each condition. **(b)** Ternary plots showing the percentage of secondary structure for each peptide (at 50 μmol L^-1^) in four different solvents: water, 60% trifluoroethanol (TFE) in water, 50% methanol (MeOH) in water, and Sodium dodecyl sulfate (SDS, 10 mmol L^-1^) in water. Secondary structure fractions were calculated using the BeStSel server^26^. Circles indicate active VEPs, while crosses represent inactive peptides.

The inactive or minimal activity of UniProtKB-2, -6, and -11 was associated with lower hydrophobicity and net charge, underscoring the important role of these parameters in facilitating membrane interactions. Conversely, ISOB-derived VEPs with enhanced normalized hydrophobicity exhibited improved antimicrobial performance. Among Conoserver-derived VEPs, an intermediate balance of hydrophobicity and net charge appeared to be optimal for activity. In Arachnoserver-derived VEPs, where all candidates were active, efficacy seemed to be driven by sequence-specific features rather than general physicochemical properties.

These findings underscore the importance of physicochemical characteristics, such as charge and hydrophobicity, in effective bacterial membrane disruption while also highlighting the significant role of sequence-specific factors in determining antimicrobial efficacy.

### Secondary structure studies

The secondary structure of short peptides is inherently dynamic, often transitioning between disordered and ordered conformations depending on the surrounding environment, particularly at hydrophobic/hydrophilic interfaces. These structural transitions are critical for defining the biological functions of peptides, including their antimicrobial activity.

To investigate the structural behavior of the synthesized VEPs, we performed Circular Dichroism (CD) spectroscopy in diverse environments: water, sodium dodecyl sulfate (SDS)/water (10 mmol LC¹), methanol (MeOH)/water (1:1, v:v), and trifluoroethanol (TFE)/water (3:2, v:v). Each medium was chosen to simulate specific physicochemical conditions relevant to peptide behavior. SDS micelles mimic biological lipid bilayers, offering a membrane-like environment conducive to evaluating interactions with bacterial membranes^19^. The TFE/water mixture is a known helical-inducer that promotes intramolecular hydrogen bonding by dehydrating peptide backbone amide groups, thereby favoring α-helical conformations^21,23^. Conversely, the MeOH/water mixture promotes interchain hydrogen bonding, stabilizing β-like structures, while hydrophobic side chains cluster to minimize contact with water, enhancing β-like conformations^25^.

CD spectra were recorded for all VEPs at 50 µmol L^-1^ over a wavelength range of 260 to 190 nm (**Supplementary Figure 8a-d**). The Beta Structure Selection (BeStSel) algorithm was employed to deconvolute the spectra and quantify the secondary structure content^26^ (**Figure 2b-e**). As expected for short peptides (<50 amino acid residues), VEPs were predominantly unstructured in water (**Figure 2b** and **Supplementary Figure 8a,e**), though with a slight propensity toward β-like conformations (f_β_ < 45%; **Supplementary Figure 8e**). A similar trend was observed in the β-inducing medium (MeOH/water), where the β-content modestly increased (**Figure 2e** and **Supplementary Figure 8d-e**).

In contrast, VEPs exhibited a pronounced structural transition in SDS micelles (**Figure 2c** and **Supplementary Figure 8c,e**) and TFE/water mixture (3:2, v:v; **Figure 2d** and **Supplementary Figure 8b,e**), adopting α-helical conformations. This shift from disordered to α-helical structures highlights their responsiveness to membrane-mimicking environments and helical-inducing media, consistent with typical behavior observed for antimicrobial peptides^6,27,28^.

Interestingly, this behavior distinguishes VEPs from other classes of encrypted peptides, including those predicted by earlier proteome mining of APEX^9^, which predominantly adopted unstructured or β-like conformations, even in membrane-like or helical-inducing environments. Similarly, small open reading frame-encoded peptides (SEPs) and bacterial proteome-derived encrypted peptides^29,30^ showed limited helical propensity under comparable conditions. Instead, VEPs exhibited a structural response more akin to archaeasins, which also demonstrated a clear transition to α-helical conformations in helical-inducing media and upon interacting with lipid bilayers^17^. These findings suggest that VEPs may be uniquely suited for membrane-associated functions, likely contributing to their observed antimicrobial efficacy.

### Mechanism of action studies

To investigate whether VEPs exert their activity via membrane-related mechanisms, we evaluated their effects on bacterial outer and cytoplasmic membranes using fluorescence assays. We used 1-(N-phenylamino)-naphthalene (NPN) assays to assess bacterial outer membrane permeabilization (**Figure 3a**). Among the peptides tested, 23 VEPs effectively permeabilized the outer membrane. Notably, Arachnoserver-18, derived from the protein M-oxotoxin-Ot2d of the spider *Oxyopes takobius*; ConoServer-6, derived from the protein Bt211 precursor, a widely studied conotoxin from the betuline cone (*Conus betulinus*); and ConoServer-7, derived from the protein Con-ins G1b precursor of *Conus geographus*, a cone snail known for having the most potent venom among the *Conus* genus^31^, showed superior permeabilization activity. Polymyxin B and levofloxacin were as controls in these experiments^11^. Overall, VEPs demonstrated permeabilization comparable to or better than other AMPs^7,32,33^ or other human- or animal-derived EPs^9,11^.

**Figure 3.**
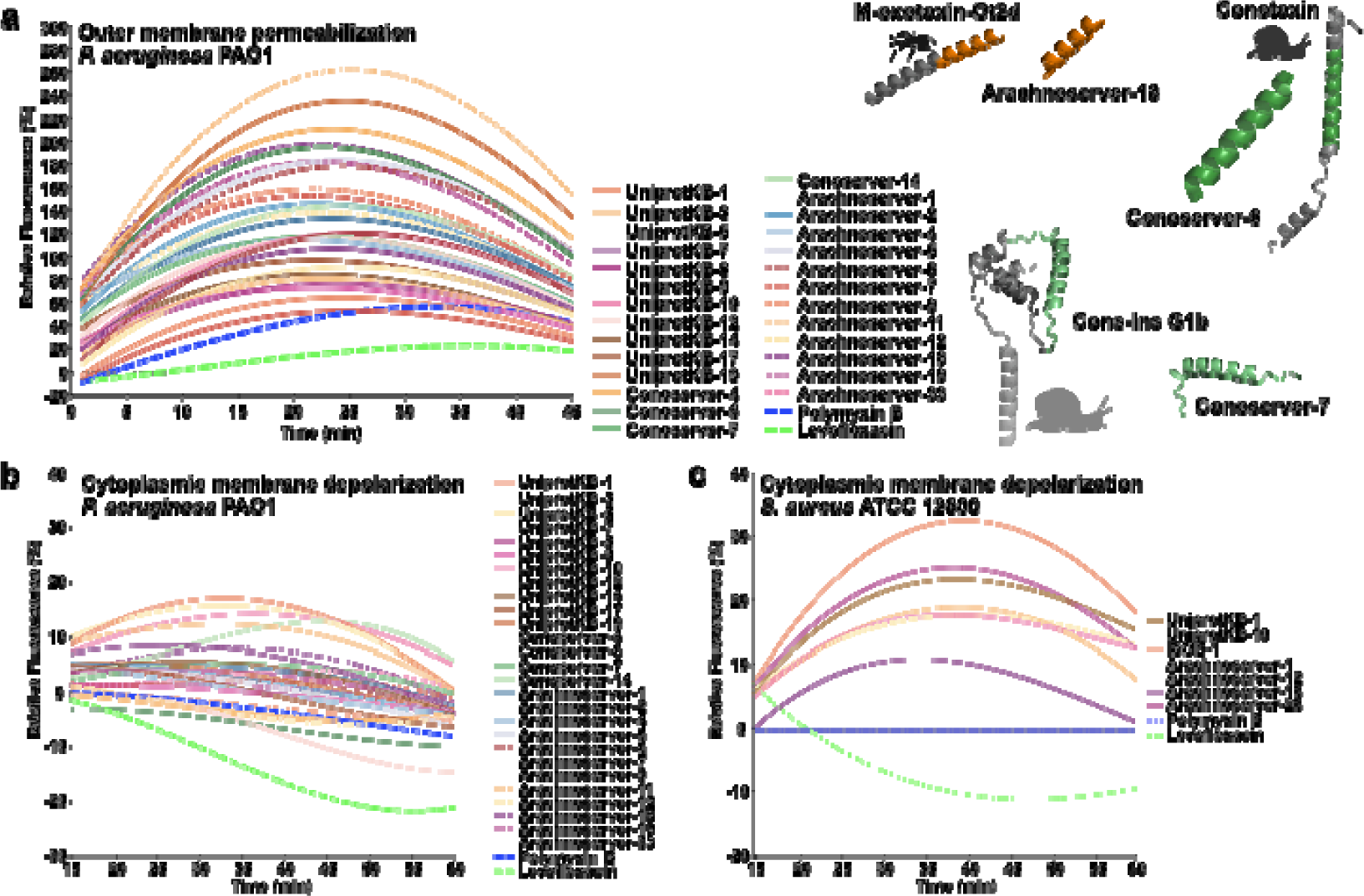
Mechanism of action of antibiotics from venoms. To assess whether VEPs act on bacterial membranes, all active peptides against *P. aeruginosa* PAO1 were subjected to outer membrane permeabilization and peptides active against *P. aeruginosa* PAO1 and *S. aureus* ATCC 12600 were tested in cytoplasmic membrane depolarization assays. The fluorescent probe 1-(N-phenylamino)naphthalene (NPN) was used to assess membrane permeabilization **(a)** induced by the tested VEPs. The fluorescent probe 3,3′-dipropylthiadicarbocyanine iodide (DiSC_3_-5) was used to evaluate membrane depolarization **(b)** caused by VEPs. The values displayed represent the relative fluorescence of both probes, with non-linear fitting compared to the baseline of the untreated control (buffer + bacteria + fluorescent dye) and benchmarked against the antibiotics polymyxin B and levofloxacin. All experiments were performed in three independent replicates. The protein and peptide structures depicted in the figure were created with PyMOL Molecular Graphics System, version 3.1 Schrödinger, LLC.

We next evaluated cytoplasmic membrane depolarization using 3,3′-dipropylthiadicarbocyanine iodide (DiSC_3_-5), a fluorophore that detects membrane potential changes. Among the 28 peptides tested against *P. aeruginosa* PAO1, 26 VEPs depolarized the cytoplasmic membrane more effectively than the control groups treated with polymyxin B and levofloxacin (**Figure 3b**)^11^. However, the depolarization efficacy of VEPs was less pronounced compared to other peptide families^29^, such as those derived from the archaeal proteome (archaeasins)^17^ and SEPs^29^. Against the Gram-positive bacterium *S. aureus*, VEPs exhibited slightly better depolarization activity than *P. aeruginosa* (**Figure 3c**), though their performance remained below that of other reported peptide depolarizers ^10,30^.

These findings suggest that VEPs primarily exert their antimicrobial effects through cytoplasmic membrane depolarization rather than outer membrane permeabilization. This mode of action aligns with that of conventional AMPs ^32,33^ and EPs^11^ but differs from certain computationally predicted peptides^29^.

### *In vitro* cytotoxicity of VEPs

Cytotoxicity was assessed using human embryonic kidney (HEK293T) cells. Some VEPs were cytotoxic at HC_50_ ≤64 µmol LC¹ (**Figure 4a**), mirroring their potent antimicrobial activity. These findings underscore the importance of fine-tuning VEP properties to balance antimicrobial efficacy with reduced cytotoxicity, guiding further peptide optimization.

**Figure 4.**
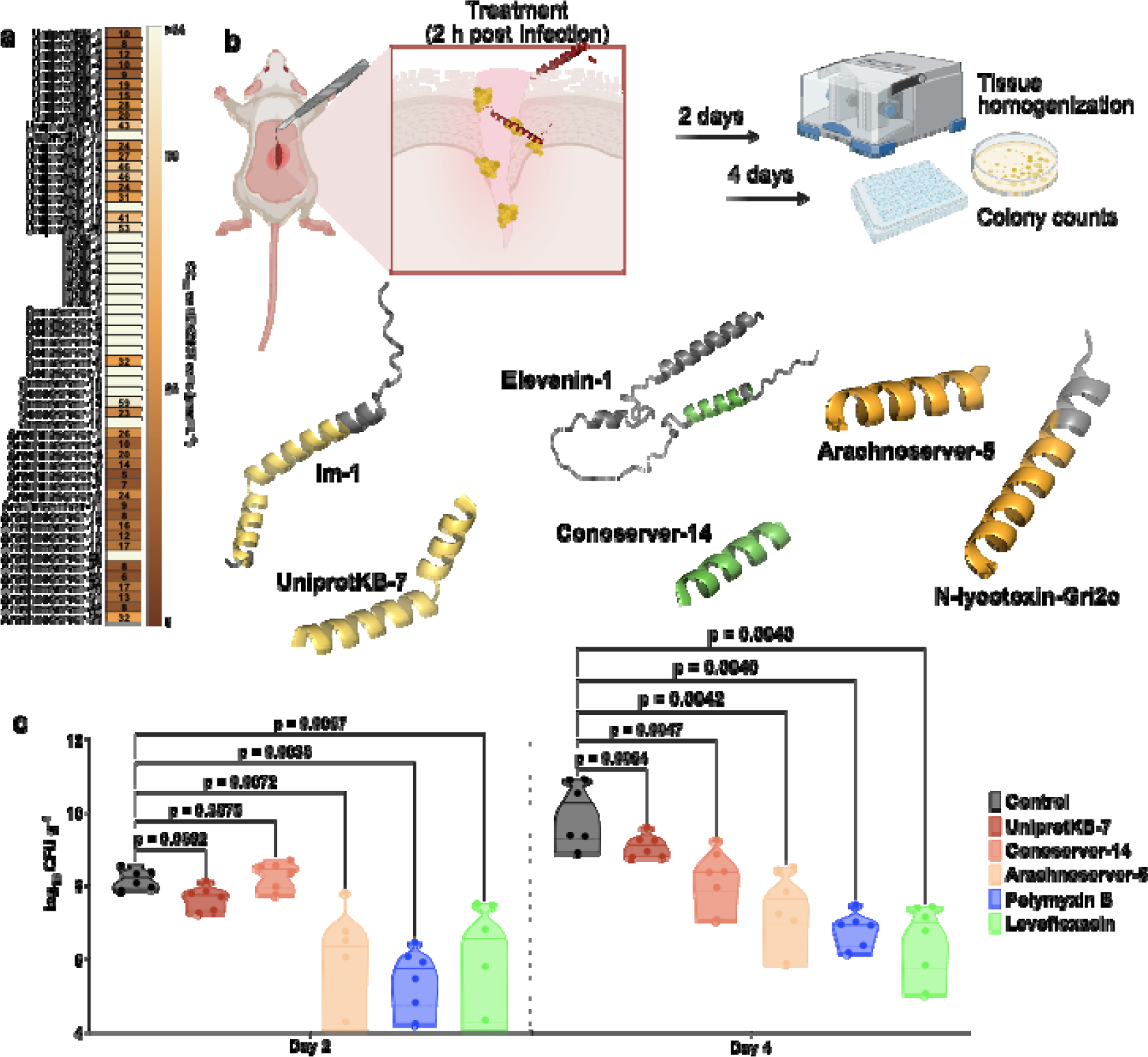
Cytotoxic and anti-infective activity of antibiotics from venoms. **(a)** Heatmap exhibiting the cytotoxic concentrations leading to 50% cell lysis (CCCC) in human embryonic kidney (HEK293T) cells, determined by interpolating dose-response data using a nonlinear regression curve. All experiments were performed in three independent replicates. **(b)** Schematic representation of the skin abscess mouse model used to assess the anti-infective activity of VEPs (nC=C6) against *A. baumannii* ATCC 19606. **(b)** UniprotKB-7, conoserver-14, and arachnoserver-5 were administered at their MIC in a single dose two hours post-infection. Arachnoserver-5 inhibited the proliferation of the infection for up to 4Cdays after treatment compared to the untreated control group at levels comparable to the control antibiotics, polymyxin B and levofloxacin.

### Anti-infective activity in preclinical animal models

To determine the *in vivo* efficacy of lead VEPs, we used a skin abscess mouse model infected with *A. baumannii*, a clinically significant pathogen (**Figure 4b**). Three VEPs demonstrated promising activity: UniProtKB-7, derived from the Im-1 toxin of the scorpion *Isometrus maculatus*; ConoServer-14, derived from the Elevenin-Vc1 protein of the cone snail *Conus quercinus*; and Arachnoserver-5, derived from the M-lycotoxin-Gri2c protein of the wolf spider *Geolycosa riograndae*.

A single topical dose of each VEP at its MIC significantly reduced bacterial counts two days post-infection. Arachnoserver-5 achieved a two-log reduction in bacterial load, comparable to the activity of polymyxin B and levofloxacin controls. Four days post-infection, all three VEPs continued to suppress bacterial growth, with Arachnoserver-5 producing a three-log reduction relative to untreated controls (**Figure 4c**). Importantly, no significant changes in body weight were observed in treated animals, indicating minimal toxicity under these conditions (**Supplementary Figure 10**).

## Discussion

This study highlights the potential of computational exploration of venom proteomes, integrating machine learning-driven predictions with experimental validation, to uncover novel antibiotic candidates. The VEPs identified in this work exhibit distinct sequence and physicochemical properties, retain membrane-active mechanisms characteristic of known antimicrobial peptides (AMPs), and demonstrate promising antimicrobial activity in both *in vitro* assays and preclinical animal models.

Our findings highlight the power of combining digital data and machine learning to accelerate antibiotic discovery. By tapping into the underexplored biodiversity of venom-derived proteins, we have uncovered a promising new class of antimicrobial agents.

### Limitations of the study

While APEX has proven effective in accelerating the discovery of novel antimicrobials, several limitations remain. One significant constraint is its reliance on discrete MIC values, which are recorded in multiples of 2, and the exclusive use of AAindex features, limiting prediction accuracy and generalizability. Additionally, the model restricts input sequence length to 50 residues, favoring peptides that are easier to chemically synthesize but compromising prediction accuracy for longer sequences. Another notable limitation is the lack of interpretability in APEX’s predictions, as it does not identify specific sequence features responsible for AMP activity, thereby limiting mechanistic insights.

To address these limitations and enhance APEX’s utility, several strategies can be implemented. Introducing self-attention mechanisms could improve model interpretability by pinpointing critical sequence features that drive antimicrobial activity. Expanding the training dataset to include longer sequences would improve prediction accuracy for peptides exceeding the 50-residue threshold. Employing data augmentation techniques could enhance generalizability across diverse peptide datasets. Additionally, integrating large language models could capture complex sequence relationships, further improving prediction accuracy and broadening APEX’s applicability.

## Acknowledgments

Cesar de la Fuente-Nunez holds a Presidential Professorship at the University of Pennsylvania and acknowledges funding from the Procter & Gamble Company, United Therapeutics, a BBRF Young Investigator Grant, the Nemirovsky Prize, Penn Health-Tech Accelerator Award, Defense Threat Reduction Agency grants HDTRA11810041 and HDTRA1-23-1-0001, and the Dean’s Innovation Fund from the Perelman School of Medicine at the University of Pennsylvania. Research reported in this publication was supported by the Langer Prize (AIChE Foundation), the NIH R35GM138201, and DTRA HDTRA1-21-1-0014. We thank Dr. Mark Goulian for kindly donating the following strains: *Escherichia coli* AIC221 [*Escherichia coli* MG1655 phnE_2::FRT (control strain for AIC222)] and *Escherichia coli* AIC222 [*Escherichia coli* MG1655 pmrA53 phnE_2::FRT (polymyxin resistant)]. We thank de la Fuente Lab members for insightful discussions. Figures created with BioRender.com are attributed as such.

Molecules were rendered using the PyMOL Molecular Graphics System, Version 3.1.1 Schrödinger, LLC.

## Author contributions

Conceptualization: MDTT, CG, CFN

Methodology: MDTT, CG, CFN

Experimental investigation: MDTT, SL

Computational investigation: CG

Visualization: MDTT, CG

Funding acquisition: CFN

Supervision: CFN

Formal analysis: MDTT, CG

Writing – original draft: MDTT, CG, CFN

Writing – review & editing: MDTT, CG, CFN

## Data availability

The main data supporting the results in this study are available within the paper and Data S1 and S2 files (DOI: 10.17632/9m4g52grhj.1). All data generated in this study, including Source Data for the figures are available from the corresponding author on reasonable request.

## Code availability

APEX is available at GitLab: https://gitlab.com/machine-biology-group-public/apex.

## Methods

### Encrypted peptides in venom proteomes

The venom protein sequences were collected from https://www.snakebd.com/ (Snakes), https://arachnoserver.qfab.org/mainMenu.html (Spider), https://www.conoserver.org/ (Carnivorous marine cone snails) and https://venomzone.expasy.org/ (Venom Zone) (access data: August 30th, 2023). 654, 2,206, 5,494 and 7,769 proteins were obtained from above four databases, respectively. Venom protein substrings ranging from 8-50 amino acids in the sequences, comprising only canonical amino acids, were considered as the venom encrypted peptides (VEPs). The venom proteins were preprocessed in three ways based on length: (1) no truncation for lengths ≥8; (2) truncation using a sliding window (range from 8 to maximum sequence length) for lengths between 8 and 50; (3) truncation using a sliding window (range from 8 to 50) for lengths >50. In total, 40,626,260 VEPs were obtained from venom proteins sequences, which were for further study.

### APEX

APEX is a bacterial strain-specific antimicrobial activity predictor^9^, and was trained on in-house peptide dataset and publicly available antimicrobial peptides (AMPs) from DBAASP^34^. Specifically, APEX is a multiple-target tasks model that can predict minimum inhibitory concentrations (MICs) values of peptides against 34 bacterial strains (*E. coli* ATCC 11775*, P. aeruginosa* PAO1*, P. aeruginosa* PA14*, S. aureus* ATCC 12600*, E. coli* AIC221*, E. coli* AIC222*, K. pneumoniae* ATCC 13883*, A. baumannii* ATCC 19606*, Akkermansia muciniphila* ATCC BAA-835*, Bacteroides fragilis* ATCC 25285*, Bacteroides vulgatus (Phocaeicola vulgatus)* ATCC 8482*, Collinsella aerofaciens* ATCC 25986*, Clostridium scindens* ATCC 35704*, Bacteroides thetaiotaomicron* ATCC 29148*, B. thetaiotaomicron* Δ*tdk* Δ*lpxF* (background: VPI 5482)*, Bacteroides uniformis* ATCC 8492*, Bacteroides eggerthi* ATCC 27754*, Clostridium spiroforme* ATCC 29900*, Parabacteroides distasonis* ATCC 8503*, Prevotella copri* DSMZ 18205*, Bacteroides ovatus* ATCC 8483*, Eubacterium rectale ATCC 33656, Clostridium symbiosum ATCC 14940, Ruminococcus obeum ATCC 29174, Ruminococcus torques ATCC 27756, methicillin-resistant S. aureus ATCC BAA-1556, vancomycin-resistant Enterococcus faecalis ATCC 700802, vancomycin-resistant E. faecium ATCC 700221, E. coli Nissle 1917, Salmonella enterica ATCC 9150 (BEIRES NR-515), S. enterica (BEIRES NR-170), S. enterica ATCC 9150 (BEIRES NR-174) and Listeria monocytogenes ATCC 19111 (BEIRES NR-106)*

### Venom encrypted peptide selection

APEX was used to predict the antimicrobial activity for the 40,626,260 encrypted peptides derived from the venom proteome. We used the mean MIC value against the eleven pathogen strains to rank and select the encrypted peptides for chemical synthesis and experimental validation. When selecting the peptides, we also make sure they met the following criteria:

1 – The selected peptide should have ≤32 μmol L^-1^ median MIC by prediction.
2 – The selected peptide should have <75% sequence similarity to all in-house peptides and publicly available AMPs.
3 – The selected peptides themselves should have <75% sequence similarity.

### Physicochemical property analysis

The twelve physicochemical properties of peptides, including normalized hydrophobic moment, normalized hydrophobicity, net charge, isoelectric point, penetration depth, tilt angle, disordered conformation propensity, linear moment, propensity to aggregation *in vitro*, angle subtended by the hydrophobic residues, amphiphilicity index, and propensity to PPII coil, were obtained from the DBAASP server^35^. Note that Eisenberg and Weiss scale^36^ was chosen as the hydrophobicity scale.

### Phylogenetic tree visualization

To obtain the phylogenetic tree, the taxon IDs of organisms obtained from four databases were uploaded to NCBI Taxonomy Common Tree (https://www.ncbi.nlm.nih.gov/Taxonomy/CommonTree/www.cmt.cgi). The resulted tree file from NCBI was then visualized via iTOL (https://itol.embl.de/).

### Peptide sequence similarity

We applied the Needleman–Wunsch algorithm in the function ‘needleall’ from the EMBOSS software package (version 6.6.0.0)^37^ to estimate the similarity between our VEP with median MIC ≤32 μmol L^-1^ and AMPs in the DBAASP dataset. The parameters used are all default, and the parameter ‘identity’ was sifted out for the graph.

### AA frequencies calculation

The function ‘ProtParam.ProteinAnalysis’ was imported from the Biopython module ‘Bio.SeqUtils.ProtParam’ (version 1.75)^38^, which was used to count the total number of amino acids in a protein sequence and calculate the percentage composition of each amino acid in a protein sequence for two levels analysis including amino acid level and sequence level.

Amino acid level:

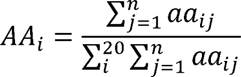

Where *aa_ij_* is the number of amino acid i in sequence j and *AA_i_* is the frequency of amino acid I. n is the total number of sequences and 20 is the total number of amino acids.

Sequence level:

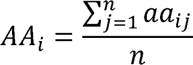

Where aa_ij_ is the frequency of amino acid i in sequence j and AA_i_ is the frequency of amino acid i. n is the total number of sequences.

### Peptide sequence space visualization

Given a peptide dataset, a similarity matrix containing the pairwise peptide sequence similarity could be calculated by previous method (Peptide sequence similarity). Uniform manifold approximation and projection (UMAP) was then used to transform the similarity matrix into a two-dimensional space. We used this space as a proxy for the peptide sequence space, and visualized the peptides’ distribution/spread/location in it.

### Peptide Synthesis

All peptides used in the experiments were purchased from AAPPTec and synthesized by solid-phase peptide synthesis using the Fmoc strategy.

### Bacterial strains and growth conditions

In this study, we used the following pathogenic bacterial strains: *Acinetobacter baumannii* ATCC 19606, *Escherichia coli* AIC221 [*Escherichia coli* MG1655 phnE_2::FRT (control strain for AIC 222)] and *Escherichia coli* AIC222 [*Escherichia coli* MG1655 pmrA53 phnE_2::FRT (polymyxin resistant; colistin-resistant strain)], *Klebsiella pneumoniae* ATCC 13883, *Pseudomonas aeruginosa* PAO1, *Pseudomonas aeruginosa* PA14, *Staphylococcus aureus* ATCC 12600, methicillin-resistant *Staphylococcus aureus* ATCC BAA-1556, vancomycin-resistant *Enterococcus faecalis* ATCC 700802, and vancomycin-resistant *Enterococcus faecium* ATCC 700221. Pseudomonas Isolation (*Pseudomonas aeruginosa* strains) agar plates were exclusively used in the case of *Pseudomonas* species. All the other pathogens were grown in Luria-Bertani (LB) broth and on LB agar. In all the experiments, bacteria were inoculated from one-isolated colony and grown overnight (16 h) in liquid medium at 37 °C. In the following day, inoculums were diluted 1:100 in fresh media and incubated at 37 °C to mid-logarithmic phase.

### Minimal inhibitory concentration assays

Broth microdilution assays were performed to determine the minimum inhibitory concentration (MIC) values of each peptide. Peptides were added to nontreated polystyrene microtiter 96-well plates and 2-fold serially diluted in sterile water from 1 to 64 μmol L^-1^. Bacterial inoculum at 2×10^6^ CFU mL^-1^ in LB or BHI medium was mixed 1:1 with the peptide. The MIC was defined as the lowest concentration of peptide able to completely inhibit the bacterial growth after 24 h of incubation at 37 °C. All assays were done in three independent replicates.

### Circular dichroism experiments

The circular dichroism experiments were conducted using a J1500 circular dichroism spectropolarimeter (Jasco) in the Biological Chemistry Resource Center (BCRC) at the University of Pennsylvania. Experiments were performed at 25 °C, the spectra graphed are an average of three accumulations obtained with a quartz cuvette with an optical path length of 1.0 mm, ranging from 260 to 190 nm at a rate of 50 nm min^-1^ and a bandwidth of 0.5 nm. The concentration of all VEPs tested was 50 μmol L^-1^, and the measurements were performed in water, mixture of water and trifluoroethanol (TFE) in a 3:2 ratio, mixture of water and methanol (MeOH) in a 1:1 ratio, and sodium dodecyl sulfate (SDS) in water at 10 mmol L^-1^, with respective baselines recorded prior to measurement. A Fourier transform filter was applied to minimize background effects. Helical fraction values were calculated using the single spectra analysis tool on the server BeStSel^26^. Ternary plots were created in https://www.ternaryplot.com/ and subsequently edited.

### Outer membrane permeabilization assays

N-phenyl-1-napthylamine (NPN) uptake assay was used to evaluate the ability of the peptides to permeabilize the bacterial outer membrane. Inocula of *P. aeruginosa* PAO1 were grown to an OD at 600 nm of 0.4 mL^-1^, centrifuged (10,000 rpm at 4 °C for 10 min), washed and resuspended in 5 mmol L^-1^ HEPES buffer (pH 7.4) containing 5 mmol L^-1^ glucose. The bacterial solution was added to a white 96-well plate (100 μL per well) together with 4 μL of NPN at 0.5 mmol L^-1^. Consequently, peptides diluted in water were added at their MIC to each well, and the fluorescence was measured at λ_ex_ = 350 nm and λ_em_ = 420 nm over time for 45 min. The relative fluorescence was calculated using the untreated control (buffer + bacteria + fluorescent dye) and polymyxin B (positive control) as baselines and the following equation was applied to reflect % of difference between the baselines and the sample:

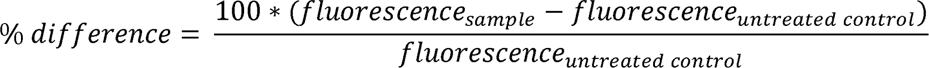

### Cytoplasmic membrane depolarization assays

The cytoplasmic membrane depolarization assay was performed using the membrane potential-sensitive dye 3,3’-dipropylthiadicarbocyanine iodide (DiSC_3_-5). *P. aeruginosa* PAO1 and *S. aureus* ATCC 12600 in the mid-logarithmic phase were washed and resuspended at 0.05 OD mL^-1^ (optical value at 600 nm) in HEPES buffer (pH 7.2) containing 20 mmol L^-1^ glucose and 0.1 mol L^-1^ KCl. DiSC_3_-5 at 20 μmol L^-1^ was added to the bacterial suspension (100 μL per well) for 15 min to stabilize the fluorescence which indicates the incorporation of the dye into the bacterial membrane, and then the peptides were mixed 1:1 with the bacteria to a final concentration corresponding to their MIC values. Membrane depolarization was then followed by reading changes in the fluorescence (λ_ex_ = 622 nm, λ_em_ = 670 nm) over time for 60 min. The relative fluorescence was calculated using the untreated control (buffer + bacteria + fluorescent dye) and polymyxin B (positive control) as baselines and the following equation was applied to reflect % of difference between the baselines and the sample:

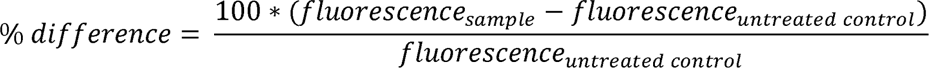

### Eukaryotic cells culture

HEK293T cells were obtained from the American Type Culture Collection (CRL-3216). The cells were cultured in high-glucose Dulbecco’s modified Eagle’s medium supplemented with 1% penicillin and streptomycin (antibiotics) and 10% fetal bovine serum and grown at 37 °C in a humidified atmosphere containing 5% CO_2_.

### Cytotoxicity assays

One day before the experiment, an aliquot of 100 μL of the cells at 50,000Ccells per mL was seeded into each well of the cell-treated 96-well plates used in the experiment (that is, 5,000 cells per well). The attached HEK293T cells were then exposed to increasing concentrations of the peptides (8–128CμmolCL^-1^) for 24 h. After the incubation period, we performed the 3-(4,5-dimethylthiazol-2-yl)-2,5-diphenyltetrazolium bromide tetrazolium reduction assay (MTT assay). The MTT reagent was dissolved at 0.5CmgCmL^-1^ in medium without phenol red and was used to replace cell culture supernatants containing the peptides (100 μL per well), and the samples were incubated for 4 h at 37C°C in a humidified atmosphere containing 5% CO_2_ yielding the insoluble formazan salt. The resulting salts were then resuspended in hydrochloric acid (0.04CmolCL^-1^) in anhydrous isopropanol and quantified by spectrophotometric measurements of absorbance at 570 nm. All assays were done as three biological replicates.

### Skin abscess infection mouse model

The back of six-week-old female CD-1 mice under anesthesia were shaved and injured with a superficial linear skin abrasion made with a needle. An aliquot of *A. baumannii* ATCC 19606 (9.6×10^5^ CFU mL^-1^; 20 μL) previously grown in LB medium until OD (optical value at 600 nm) 0.5 and then washed twice with sterile PBS (pH 7.4, 10,000 rpm for 2 min) was added to the scratched area. Peptides diluted in sterile water at MIC value were administered to the wound area 2 h after the infection. Two- and four-days post-infection animals were euthanized, and the scarified skin was excised, homogenized using a bead beater (25 Hz for 20 min), 10-fold serially diluted, and plated on McConkey agar plates for CFU quantification. The experiments were performed using six mice per group (n = 6). The skin abscess infection mouse model was revised and approved by the University Laboratory Animal Resources (ULAR) from the University of Pennsylvania (Protocol 806763).

## Quantification and statistical analysis

### Reproducibility of the experimental assays

Unless otherwise stated, all assays were performed in three independent biological replicates as indicated in each figure legend and Experimental Models and Methods details sections. The values obtained for cytotoxic activity were estimated by non-linear regression based on the screen of peptides in a gradient of concentrations and represent the cytotoxic concentration values needed to lyse and kill 50% of the cells present in the experiment. In the skin abscess mouse model, we used six mice per group following established protocols approved by the University Laboratory of Animal Resources (ULAR) of the University of Pennsylvania.

### Statistical tests

In the mouse experiments, all the raw data underwent log_10_ transformation and the statistical significance was determined using one-way ANOVA followed by Dunnett’s test. All the P-values are shown for each of the groups, all groups were compared to the untreated control group.

### Statistical analysis

All calculation and statistical analyses of the experimental data were conducted using GraphPad Prism v.10.0.2. Statistical significance between different groups was calculated using the tests indicated in each figure legend. No statistical methods were used to predetermine sample size.

## Supplementary Information

**Supplementary Table 1.** Database-sourced venom protein and VEP candidates.

| Database | Number of<br>proteins mined | Number of candidate antibiotics<br>(MIC $\leq 32 \mu\text{mol L}^{-1}$) | Number of candidates removed<br>by similarity to known AMPs | Candidates<br>(diversity filter) |
| --- | --- | --- | --- | --- |
| ConoServer | 5494 | 377 | 377 | 26 |
| ArachnoServer | 2206 | 2205 | 2154 | 80 |
| ISOB | 654 | 179 | 40 | 7 |
| VenomZone<br>(UniProtPK) | 7769 | 4618 | 3731 | 273 |

**Supplementary Table 2.**
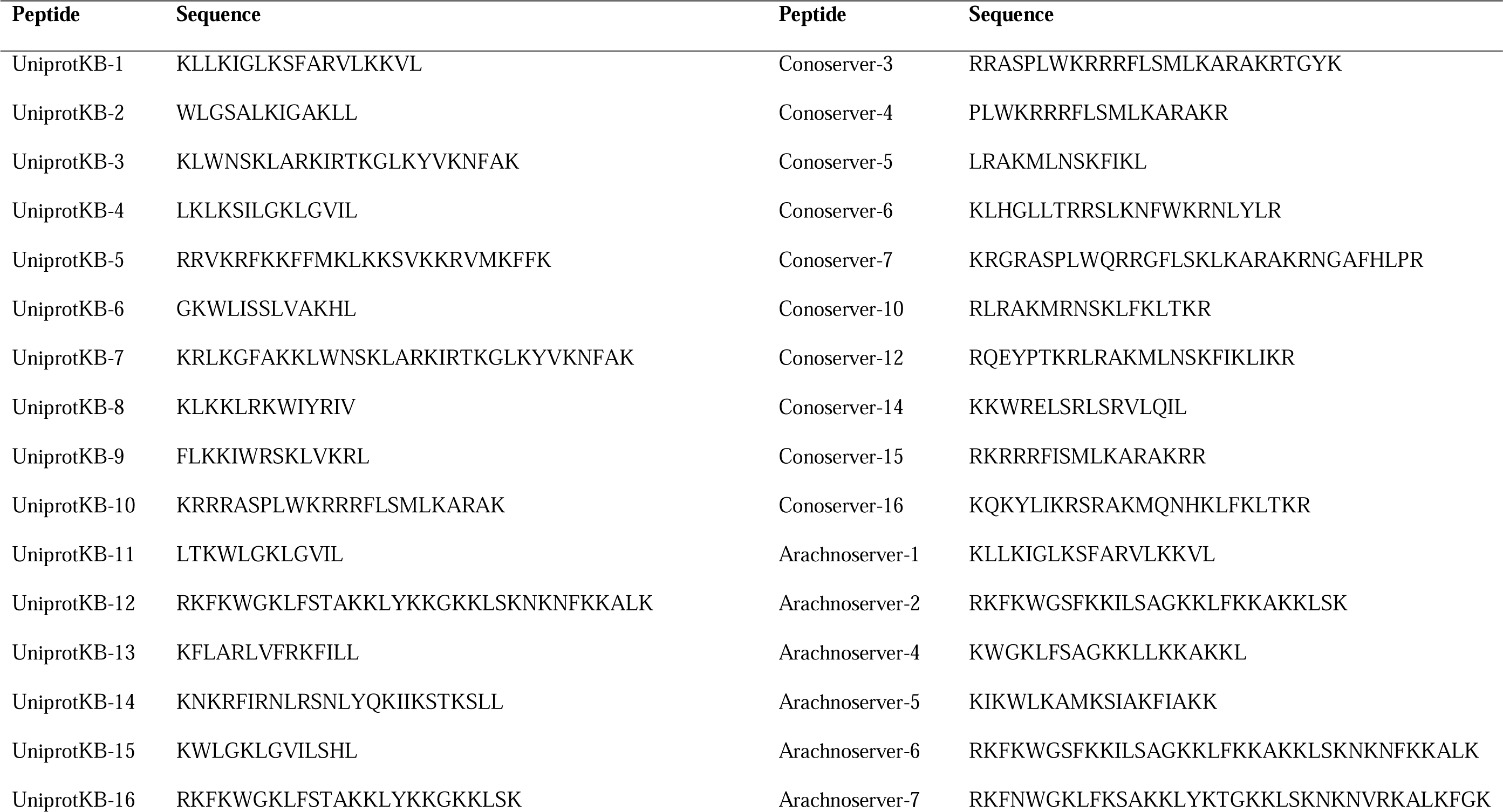
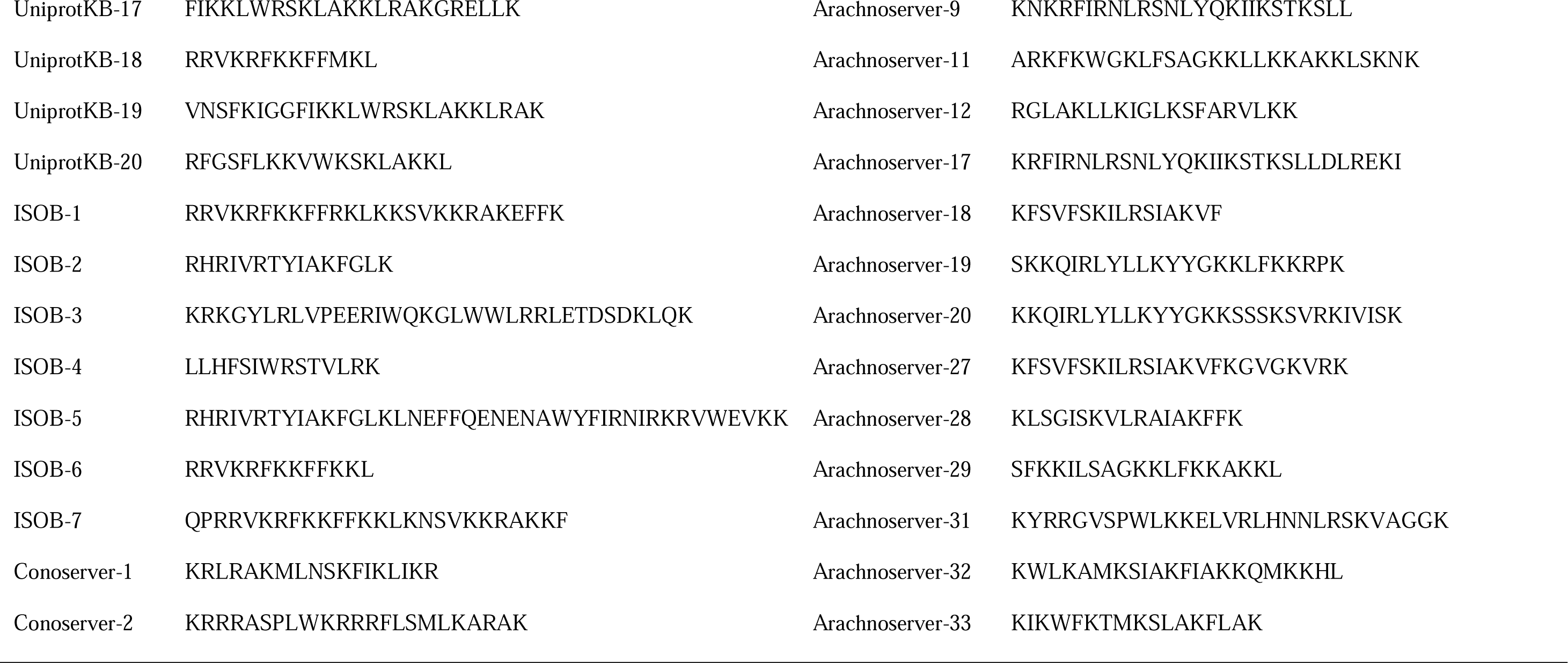
Antibiotics selected for synthesis and experimental validation.

**Supplementary Figure 1.**
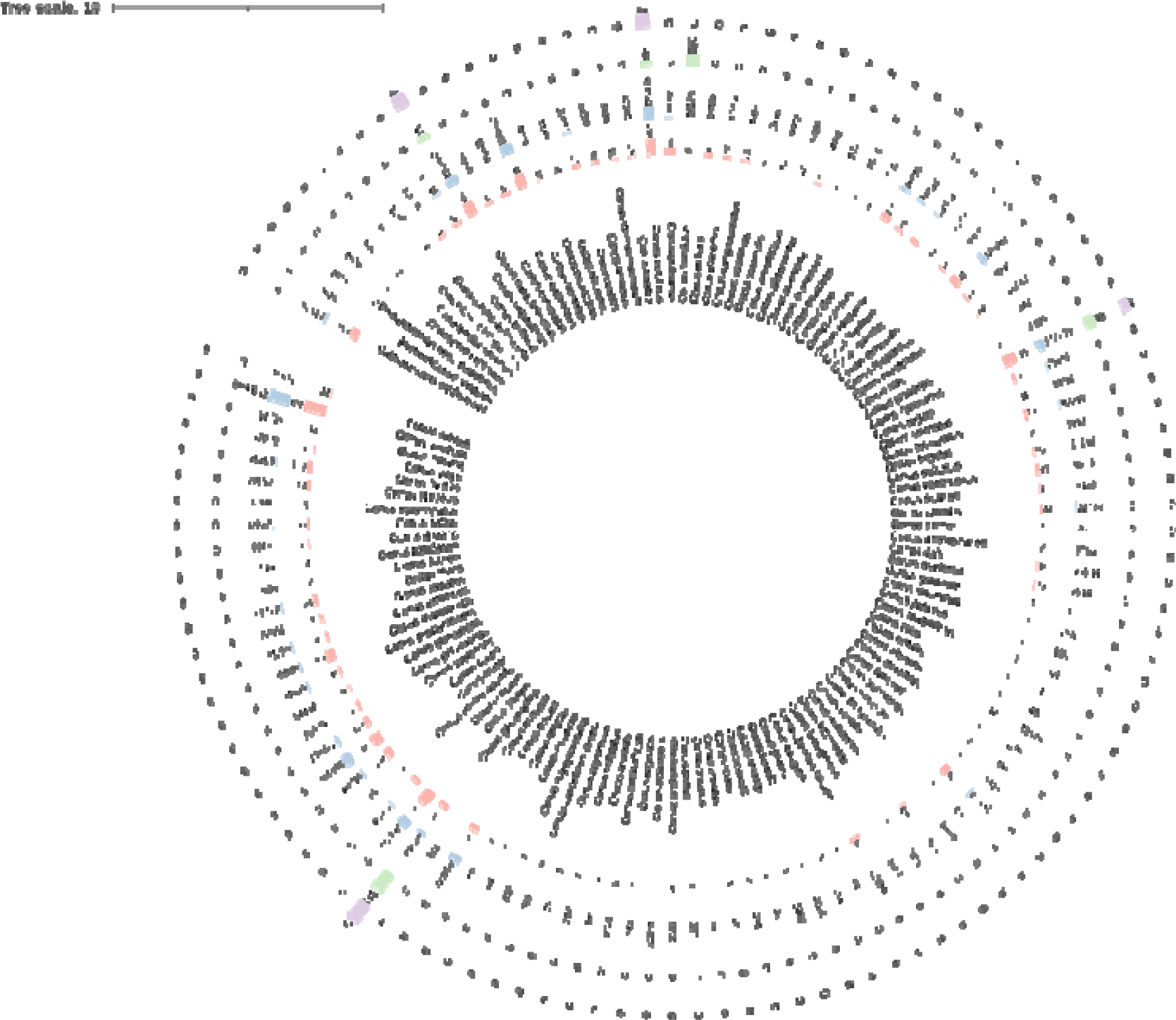
Phylogenetic analysis of venom protein, peptide, predicted AMP, and verified AMP across species in ConoServer. This phylogenetic tree illustrates the distribution of venom proteins, peptides, predicted antimicrobial peptides (AMPs), and experimentally verified AMPs among species in ConoServer. The tree was constructed using taxon IDs of organisms. From the inside to the outside, circle 1: Venom protein count per organism; Circle 2: Peptide count derived from venom proteins per organism; Circle 3: Predicted AMP count from venom proteins per organism; Circle 4: Experimentally verified AMP count per organism.

**Supplementary Figure 2.**
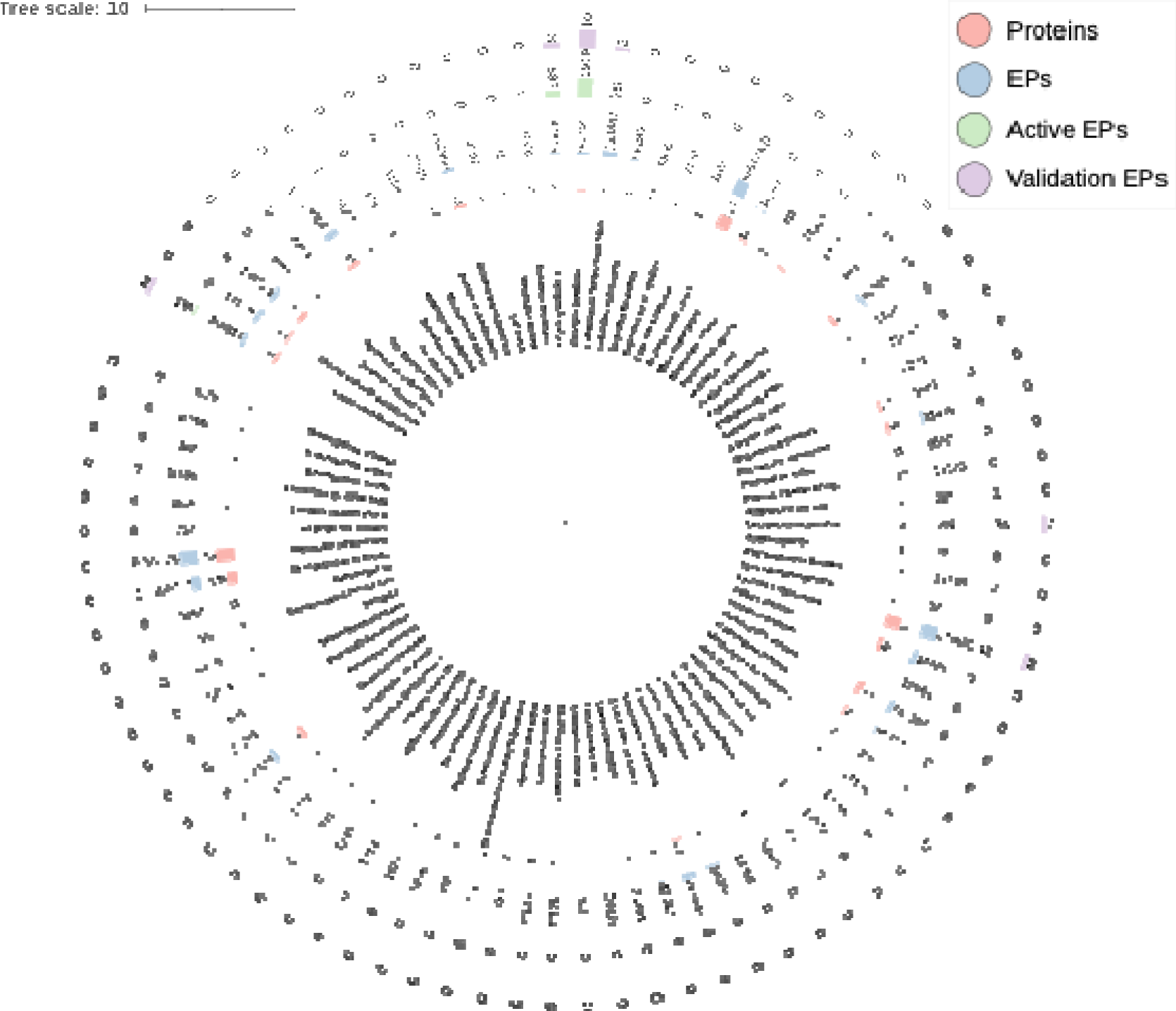
Distribution analysis of venom protein, peptide, predicted AMP, and verified AMP across species in ArachnoServer. This phylogenetic tree illustrates relationship distance of different organisms. The tree was constructed using taxon IDs of organisms. From the inside to the outside, circle 1: Venom protein number; Circle 2: Peptide number; Circle 3: Predicted AMP number; Circle 4: Experimentally verified AMP number.

**Supplementary Figure 3.**
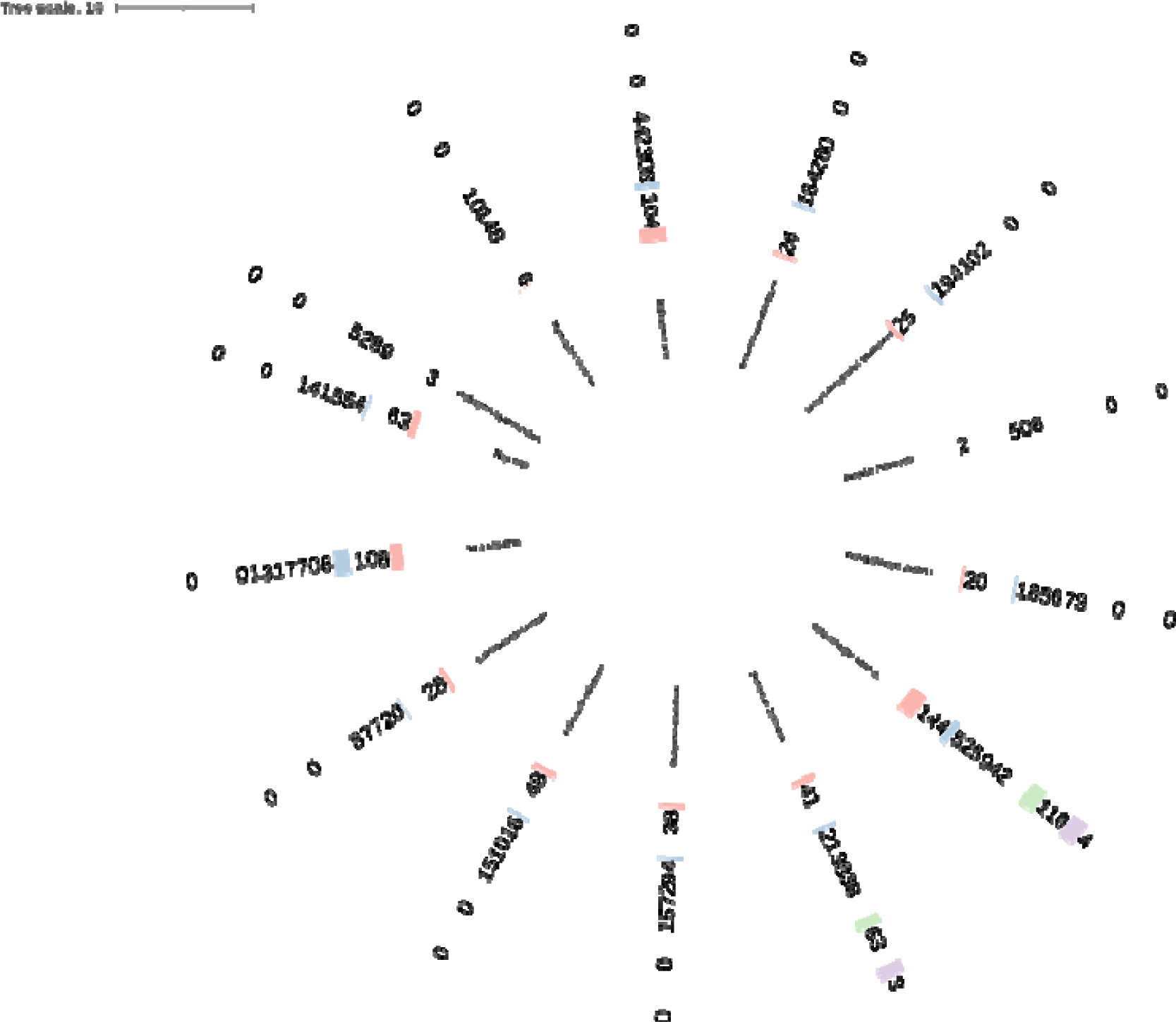
The distribution of venom protein, peptide, predicted AMP, and verified AMP across species in ISOB. This phylogenetic tree illustrates the phylogenetic relationship of species in ConoServer, which contained the number of venom proteins, peptides, predicted antimicrobial peptides (AMPs), and experimentally verified AMPs among. The four circles represent, from the inside to the outside, number of venom protein, number of peptides, number of predicted AMP, number of experimentally validation AMP.

**Supplementary Figure 4.**
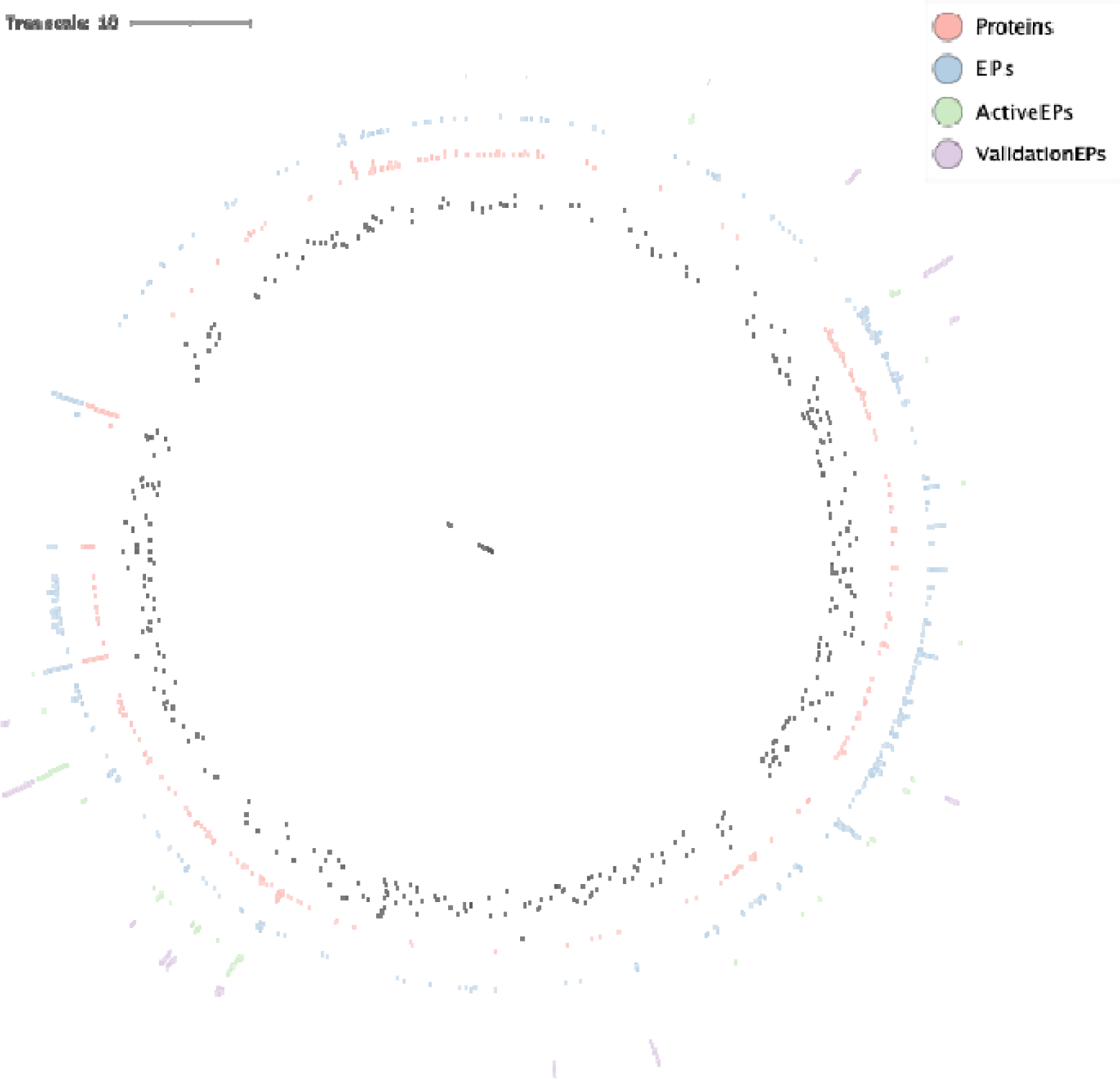
Phylogenetic analysis of species in UniProt and distribution analysis of the number of venom protein, peptide, predicted AMP, and verified AMP across species in UniProt. This evolutionary tree shows the evolutionary relationships between species in ConoServer. The four circles represent, from the inside to the outside, the number of venom proteins contained in each organism, the number of peptides produced by the venom proteins of each organism, the number of predicted AMPs contained in each organism, and the number of experimentally verified AMPs contained in each organism.

**Supplementary Figure 5.**
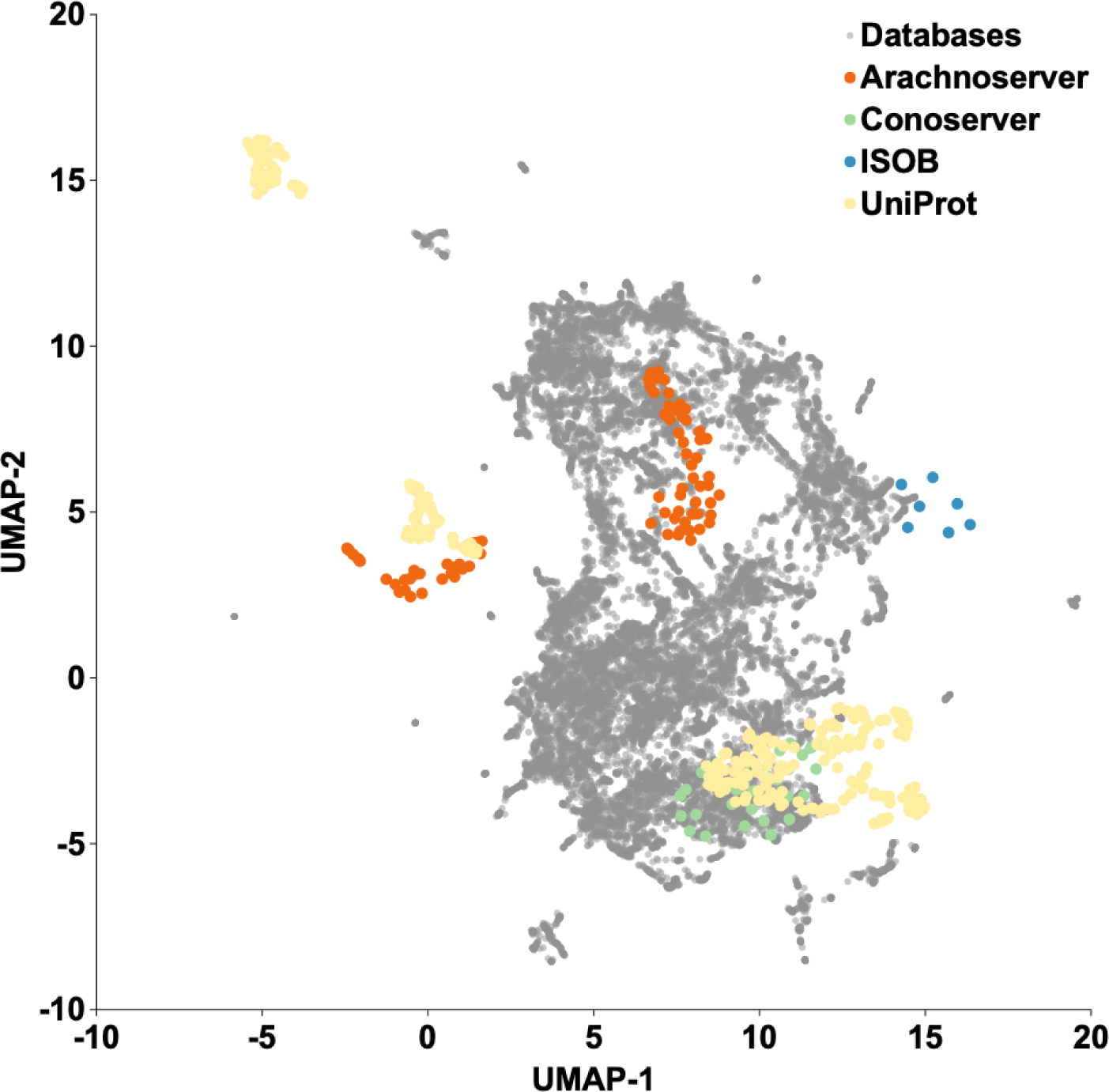
Physicochemical properties space exploration using a similarity matrix. The graph illustrates a bidimensional physicochemical properties space visualization of peptide sequences found in DBAASP and antimicrobial venom-derived EPs (VEPs) discovered by APEX in venom proteins from multiple source organisms. Sequence alignment was used to generate a similarity matrix for all peptide sequences in DBAASP and the predicted antimicrobial VEPs (see also **Data S1**). Each row in the matrix represents a feature representation of a peptide based on its amino acid composition. Uniform Manifold Approximation and Projection (UMAP) was applied to reduce the feature representation to two dimensions for visualization.

**Supplementary Figure 6.**
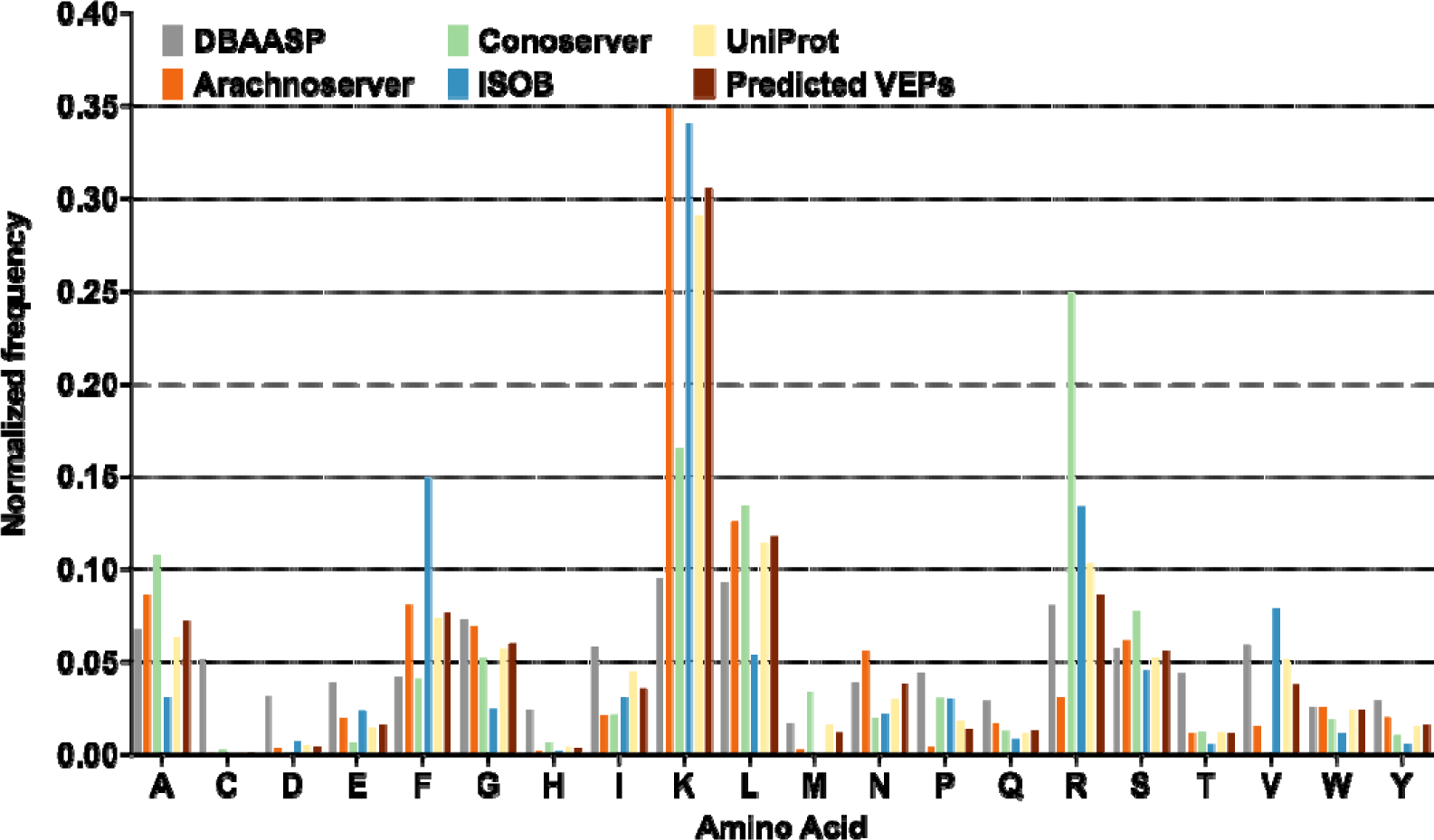
VEPs amino acid frequency at amino acid residues level. Comparison between VEPs and known antimicrobial peptides (AMPs) from the DBAASP, APD3, and DRAMP 3.0 databases at amino acid level.

**Supplementary Figure 7.**
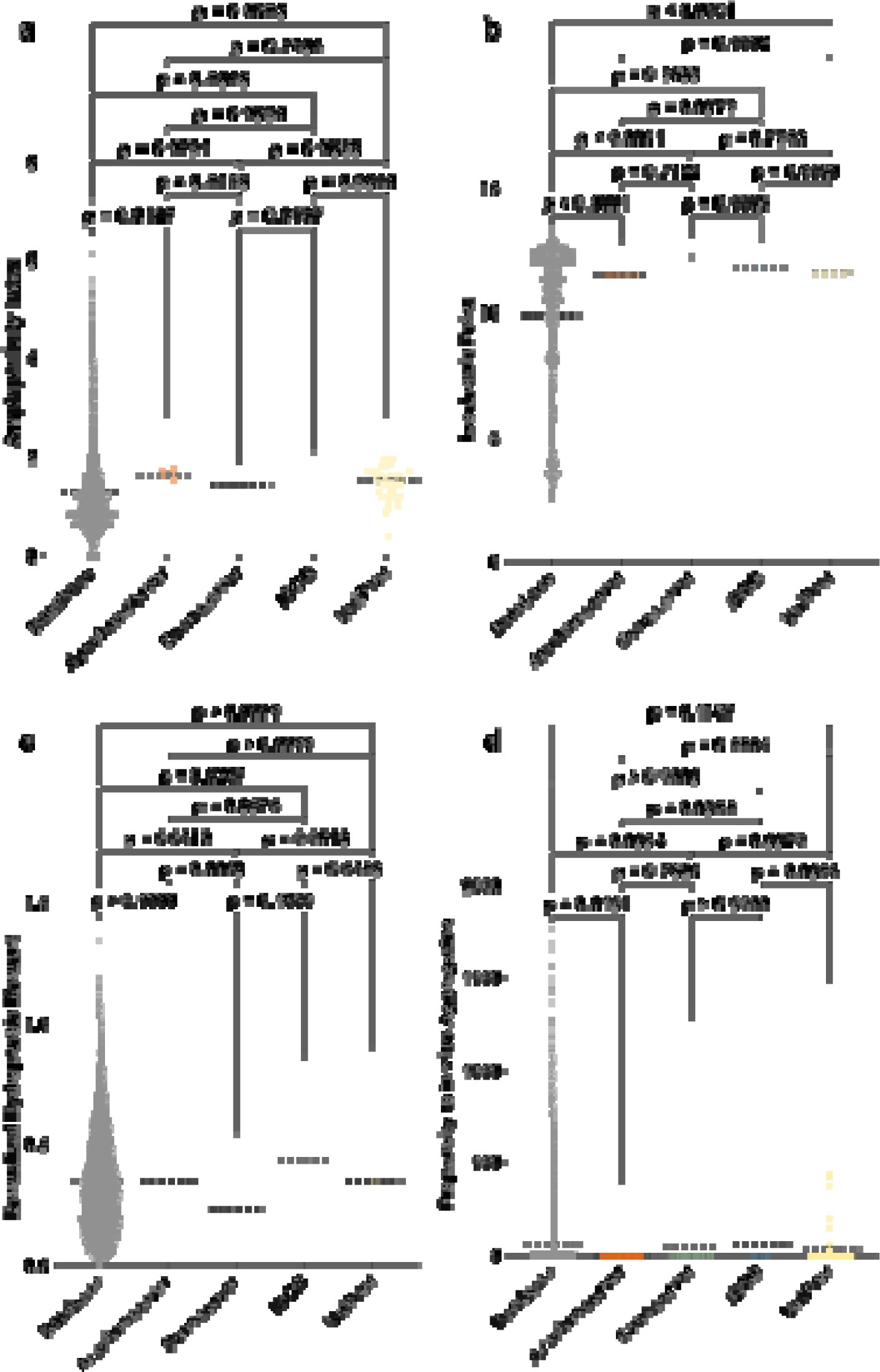
Physicochemical features of VEPs compared to AMPs from databases (DBAASP, APD3, and DRAMP 3.0). **(a)** Amphiphilicity Index, **(b)** Isoelectric Point, and **(c)** Hydrophobic moment normalized by peptide length, reflecting the amphipathicity of the molecules, which directly influences their interactions with bacterial membranes. **(d)** Propensity to aggregate *in vitro*, correlated with the supramolecular arrangement of the molecules and potential toxicity. Statistical significance was determined using two-tailed t-tests followed by the Mann-Whitney test; p values are shown in the graph. The solid line within each box represents the mean value for each group.

**Supplementary Figure 8.**
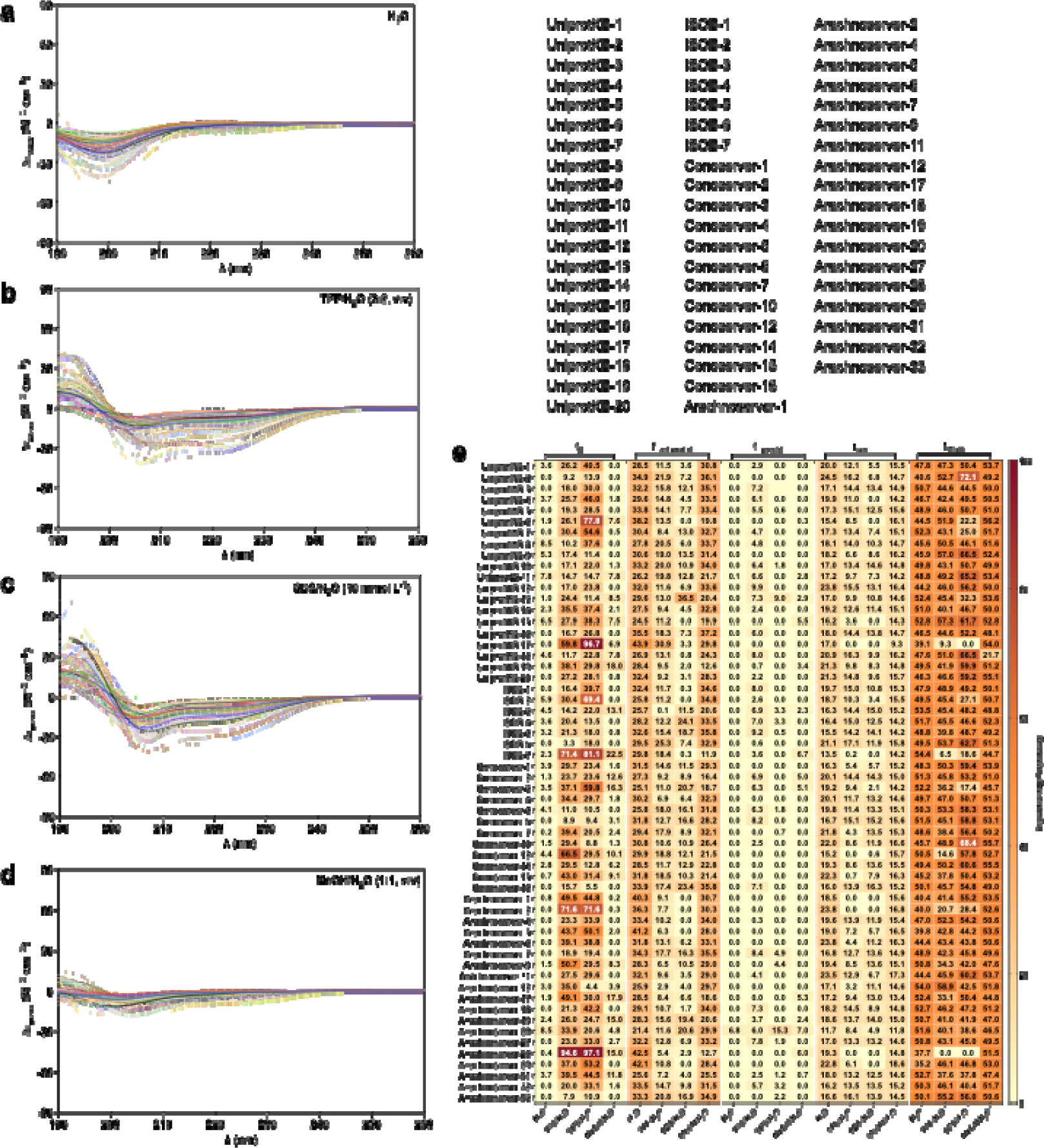
Circular dichroism spectra of VEPs. Circular dichroism experiments were conducted with peptides from venoms using a J-1500 Jasco circular dichroism spectrophotometer. The spectra were recorded in four different media: **(a)** water, **(b)** 60% trifluoroethanol in water, and **(c)** sodium dodecyl sulfate (SDS) in water (10 mmol L^-1^), and **(d)** 50% methanol in water, after three accumulations at 25 °C, using a 1mm path length quartz cell, between 260 and 190 nm at 50 nm min^-1^, with a bandwidth of 0.5 nm. The concentration of all peptides tested was 50 μmol L^-1^. **(d)** Heatmap with the percentage of secondary structure found for each peptide in the four different solvents. Secondary structure fraction was calculated using the BeStSel server^15^.

**Supplementary Figure 9.**
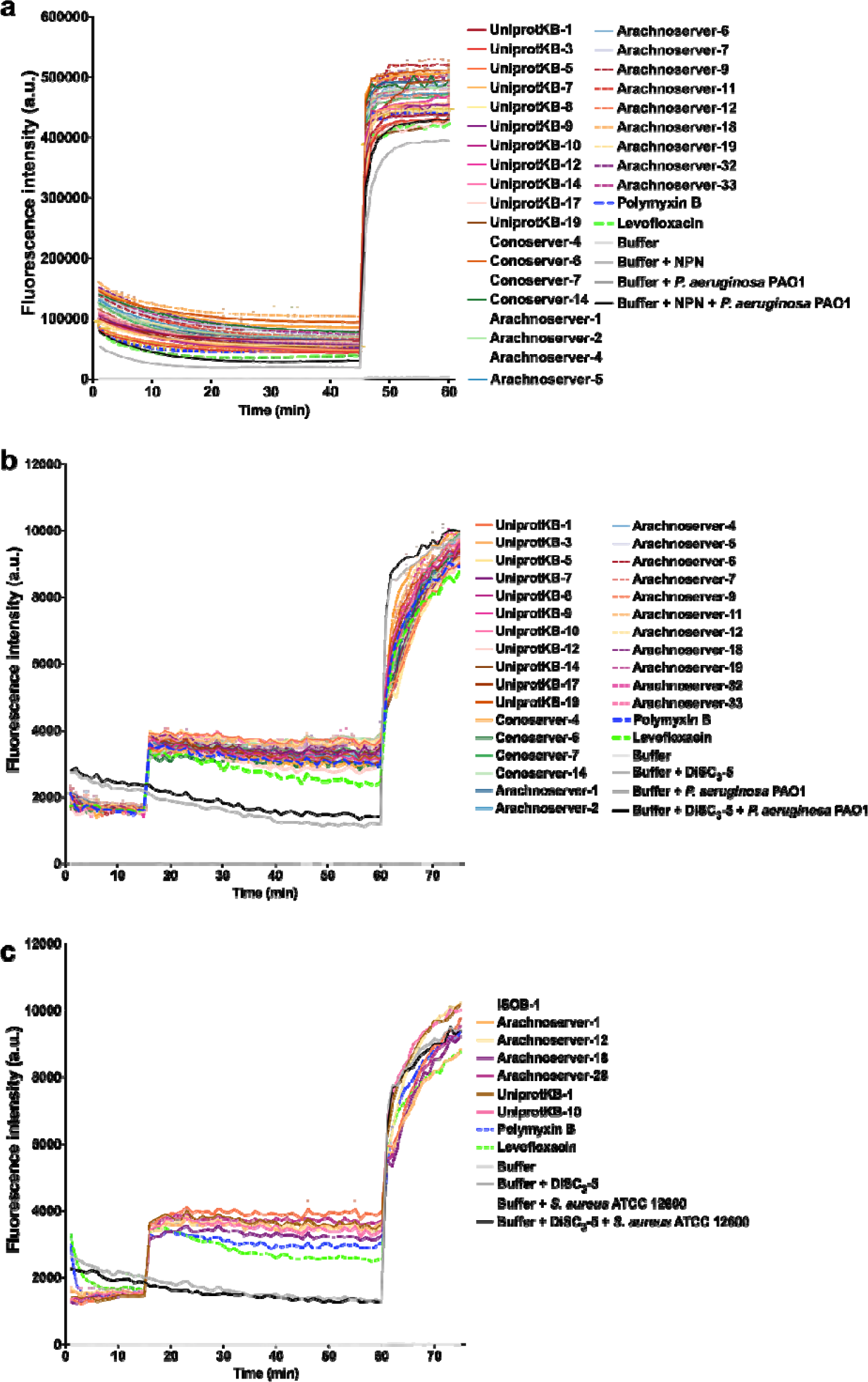
Outer membrane permeabilization and cytoplasmic membrane depolarization of *P.aeruginosa* PAO1 and *S. aureus* ATCC 12600 induced by VEPs. **(a)** Outer membrane permeabilization was assessed using the probe 1-(N-phenylamino)naphthalene (NPN), showing the permeabilization effects of VEPs active against *P. aeruginosa* PAO1. **(b)** Membrane depolarization assays were performed using the hydrophobic probe 3,3′-dipropylthiadicarbocyanine iodide (DiSC_3_-5) on all VEPs active against *P. aeruginosa* PAO1 and *S. aureus* ATCC 12600. Polymyxin B and levofloxacin served as antibiotic controls, while buffer, buffer with the probe, and buffer with both probe and bacteria were used as baseline controls for fluorescence. The panels display the raw fluorescence intensity data obtained from the experiments. Error bars are the standard deviation obtained from the three replicates.

**Supplementary Figure 10.**
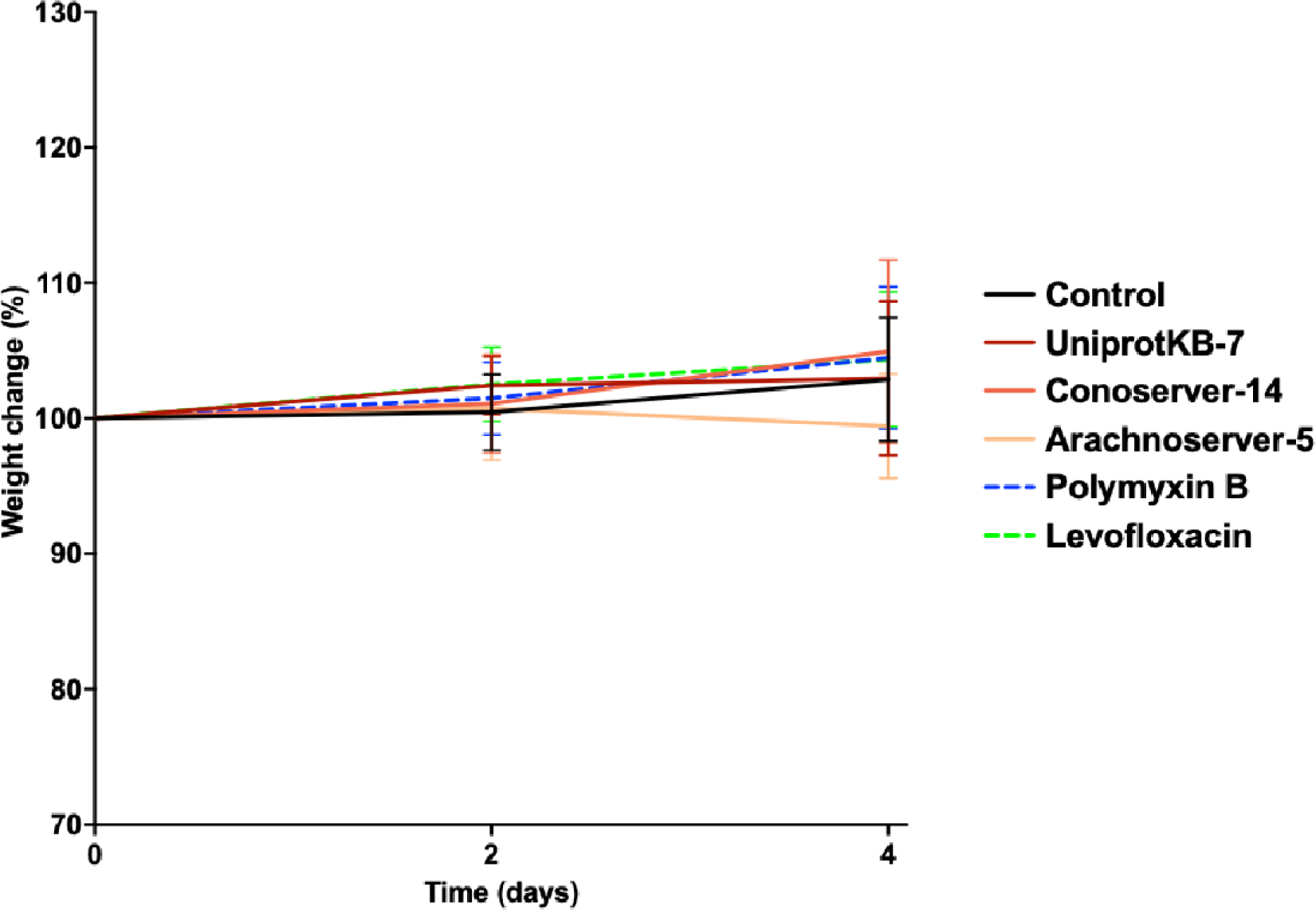
Weight change monitoring in skin abscess mouse model infected with *A. baumannii*. Mouse weight was monitored throughout the duration of the skin abscess model (4 days total) to assess potential toxic effects of both the bacterial load and the VEPs.

